# The Flexible, Versatile and Fast GreGT Platform of Fully Deleted Helper-Virus Independent Adenoviral Vectors

**DOI:** 10.1101/2024.04.04.588156

**Authors:** Yan Qi, Xianghua Zhang, Janae Wheeler Cull, Charles Wall, William Maslanik, Uwe D. Staerz

**Author notes:** Correspondence: Uwe D. Staerz, Greffex, Inc., 12635 East Montview Blvd., Aurora, CO, 80045.

## Abstract

Gene transfer (GT) vectors have diverse applications. They have been used to restore cellular activities by reconstituting normal cellular functions, by delivering therapeutic compounds, and by priming immune responses as genetic vaccines. Viruses, nature’s gene delivery vehicles, have formed the basis of most GT vectors. The biology of the underlying virus may, however, hamper their use. For instance, retroviral vectors with their ability to integrate into the genome, may cause malignant transformations. Vectors designed as minor variants of the relatively benign, yet complex adenovirus (Ad) excite vigorous immune responses, thus limiting their therapeutic effects. Deleting Ad vectors of all endogenous Ad genes brought their beneficial features to the front, such as their ability to transduce cells of many types with high efficiency and to deliver large genetic payloads. Earlier production schemes of fully deleted Ad (fdAd) vectors depended on helper virus constructs to deliver the vector production information *in trans*. They suffered from contaminations with the helper virus or the recombination of replication competent adenoviruses (RCA). We previously developed a novel transfection-based helper virus-independent Ad vector encapsidation technology that avoided these pitfalls. It was built on a vector genome and a vector packaging module. We have now optimized our approach into the GreGT *plug-and-play* platform so that a new vector can be delivered in about four weeks. The GreGT system is built upon a set of base vector genome modules that can be quickly loaded with a new application, and a set of packaging modules that allow their encapsidation into capsids of different Ad serotypes. As both components can be freely combined, the GreGT platform is endowed with high degrees of flexibility and versatility. Finally, the deletion of all endogenous Ad genes from the vector genome limits the interference by anti-Ad immune responses. It also increases the genetic payload capacity to levels unique to this system.

## INTRODUCTION

Engineered viruses as basis of gene therapy (GT) were first envisioned as early as 1970 (1). They found their first major clinical success in the treatment of an X-linked form of severe combined immune deficiency (SCID) (2). For these trials, a Moloney murine leukemia-derived vector was engineered to deliver the common alpha-chain of cytokine receptors to hemopoietic stem cells (3). Long-term reconstitution of immune functions was observed for most participants. Yet, these clinical trials illustrated the dangers inherent in a vector based on a retrovirus. About 30% of recipients developed life-threatening acute lymphatic leukemias (4,5). Another GT vector, albeit built on a benign virus, nevertheless proved problematic. In a clinical trial, an early generation Ad (egAd) vector minimally modified by removal of the E1 Ad gene was used to deliver the ornithine transcarbamylase (OTC) gene to patients suffering from OTC deficiency (6,7). One patient succumbed to severe liver complications most probably caused by an explosive innate immune response (8). Specific immune responses can also interfere with the GT success, especially when vectors are deployed that carry numerous endogenous viral genes (9,10). These observations reveal that besides the choice of the underlying agent, minor manipulations of the virus may not be sufficient to convert even a relatively safe virus, such as an Ad, into a suitable GT vector.

As exemplified by Ad vectors, a crucial step in reducing the non-therapeutic activity and at the same time, the immune target size of a GT vector, is the removal of as many, optimally all endogenous viral genes from an egAd vector (11). It was hypothesized that Ad vectors will then realize their full potential. The immune responses and toxicity caused by the endogenous Ad genes would be limited. Their genetic stability, large payload, and their broad infectivity would move to the forefront. Ad vectors *gutted* of all Ad genes can only be packaged when the necessary genetic programs are provided *in trans*. In one approach, this was achieved with a helper virus, whose encapsidation was minimized by excising its packaging site during vector replication (12). However, this process failed to prevent the contamination of Ad products with the non-therapeutic helper virus (13). Another strategy employed a baculovirus-adenovirus hybrid construct to deliver the packaging machinery. In this case, scale-up production resulted in the appearance of significant numbers of RCAs (14). We developed a new Ad vector production technology, named GreGT, in which the genetic program necessary to encapsidate a fully deleted Ad vector genome was provided by a circular packaging expression plasmid (15). This approach avoided the use of a helper virus and prevented the emergence of RCAs (*see below*). The validity of the idea was subsequently confirmed by others (16). We show here that we have further optimized GreGT into a flexible and versatile *plug-and-play* vector platform that delivers new applications in less than a month, allows diverse combinations of the vector genome and capsid, limits anti-Ad immune responses, and provides space for large transgene constructs.

## RESULTS AND DISCUSSION

### Issues

Initially we used the egAd technology to produce gene transfer vectors for different applications. In one example, an egAd vector, called edAdCD8, was adapted to express the CD8 α-chain (egAd8). In a murine islet transplantation model, we were able to demonstrate that tissues transduced to express the CD8 α-chain were protected from rejection when grafted into fully allogeneic recipients (17). As no supportive immune suppressive therapy was needed, immune responses, except those directed to the graft, remained normal. Additional experiments supported our notion that the egAdCD8 vector could similarly support the long-term acceptance of other tissues (*data not shown*). However, these experiments highlighted important shortcomings of an egAd-based vector. The egAdCD8 vector similar to other egAd vectors, exhibited a small therapeutic index. When we transduced the fragile pancreatic islets *in vitro*, we found that the egAdCD8 exhibited significant toxicity, once a certain multiplicity-of-infection (MOI) levels were surpassed (*data not shown*). Our studies with egAd vectors also confirmed that strong immune responses against the carried were raised against the vector carrier (9,18). When we injected Balb/c mice via the *intra muscular* (*i.m.*) route with an Ad vector that did not carry a transgene expression cassette (Ad(empty)), potent humoral immune responses against the Ad vector carrier were raised after a single dose (**Figure 1**). Others had also found that the deletion of all endogenous Ad genes from an Ad vector reduced its immunogenicity of an Ad vector and in parallel enhanced its ability to provide efficient and long-lasting gene therapy (19–23). The first systems designed to package fdAd relied on different helper virus constructs to deliver the packaging information *in trans.* (11,12). They suffered from issues of helper virus and RCA contaminations (12,13,14). We therefore decided to embark on the development of new fdAd technology (15), whose optimization is the topic of the present manuscript.

**Figure 1:**
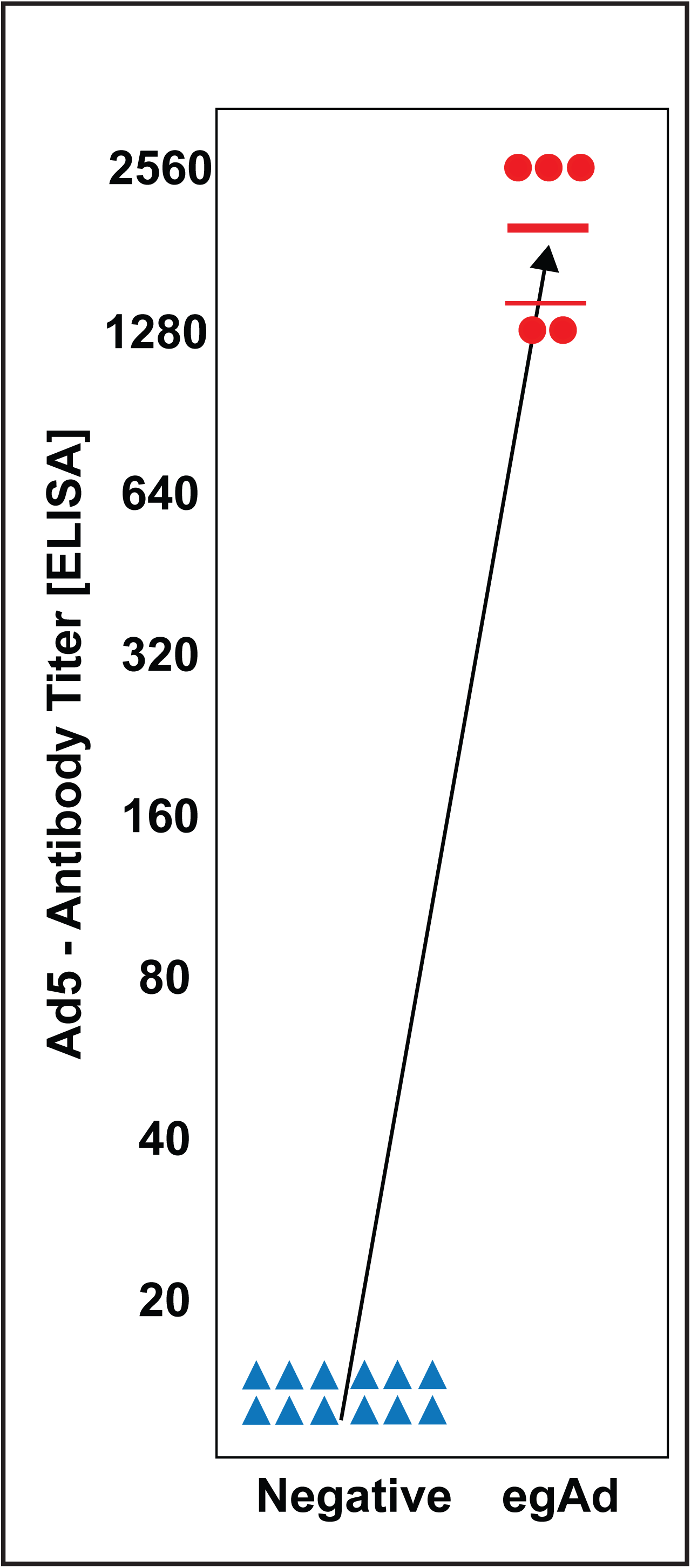
Capsid Immune Responses: Sera of negative control mice (blue triangle) that had not been exposed to an Adenoviral vector and of test mice (red filled circles) that had received an *intra muscular* injection of the egAd(empty) vector were tested for the presence of serum antibodies reactive with Adenoviral vector proteins.

### Platform Components

Our optimized fdAd technology platform, named GreGT, is built onto two genetic independent components and an engineered host cell. It consists of a set of base **genome modules** that are deleted of all Ad genes, different circular **packaging plasmids** that deliver Ad late genes *in trans*, and a **packaging cell** that has been improved for enhanced vector production. An overview of our approach is depicted in **Figure 2**.

**Figure 2:**
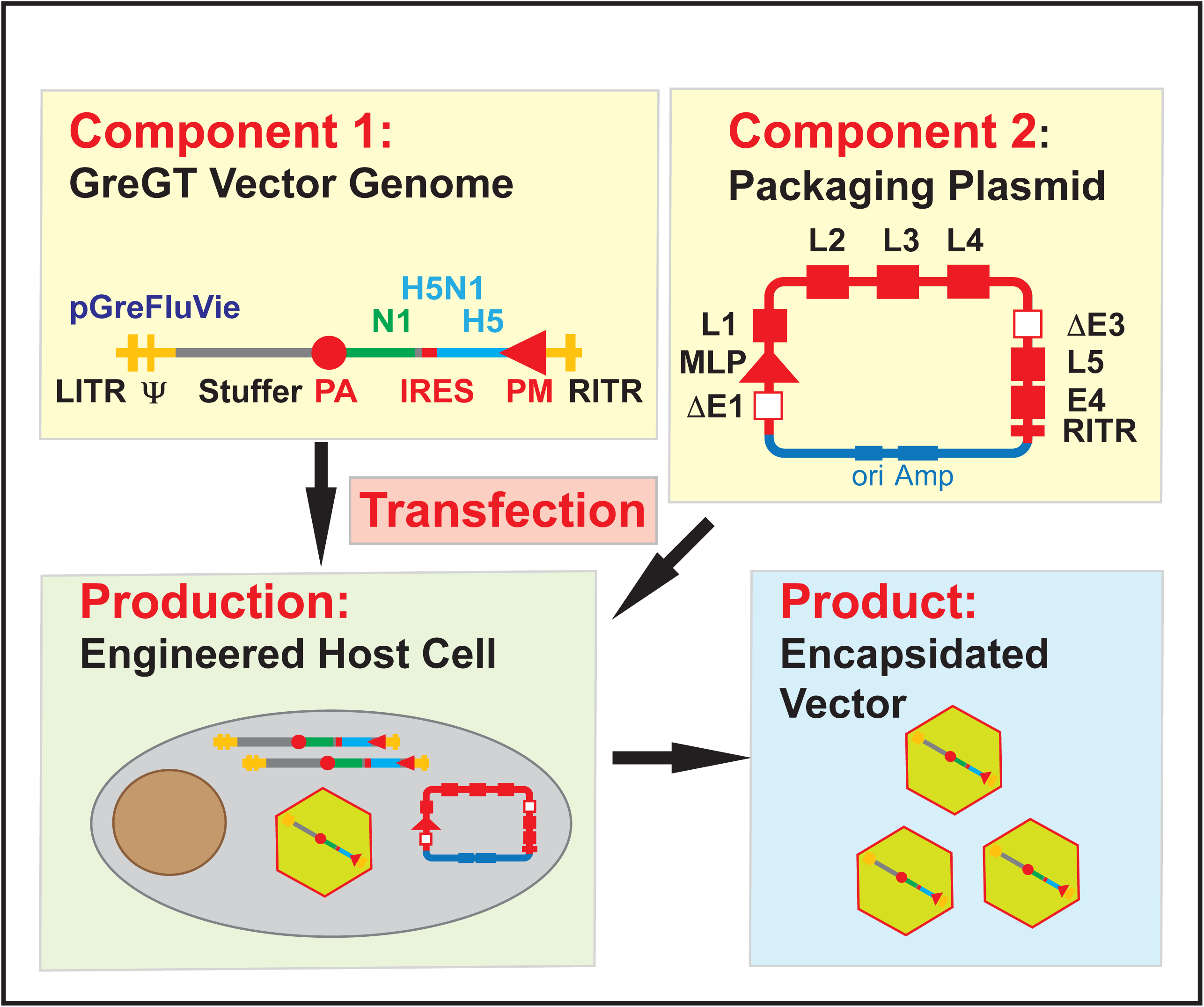
The Design of the GreGT Vector Platform: The GreGT vector platform uses the co-transfection of a linearized GreGT vector genome (Component 1) and a circular packaging expression plasmid (Component 2) into Engineered Host Cells (Production) to produce fully deleted Adenoviral vectors (Product).

#### Fully Deleted Ad Vector Modules and Genomes

The basic designs of the different vector genome plasmids were previously present in several patent applications (15). The plasmids (GreS, GreM, GreL, and GreV) were constructed to accept transgene constructs of different lengths of 1 to 8 kb (GreS), 9 to 17 kb (GreM), 18 to 25 kb (GreL) and 26 to 33 kb (GreV). They carried the left ITR (LITR), the right ITR (RITR), and a packaging signal (Ψ) derived from the human Ad serotype 5 (**Figure 3A**), but none of the endogenous Ad genes. A multiple cloning site (MCS) was fitted between a CMV immediate early promoter/enhancer and a human growth hormone polyadenylation site. Once a transgene has been moved into one of the base vector modules, a complementary *stuffer* fragment derived form the human housekeeping gene 5-aminoimidazole-4-carboxamide ribonucleotide formyltransferase gene (ATIC) is added using complimentary *recombineering* (**Figure 3B**). It compensates for the deleted Ad genes and completes the genome construct to an efficiently packageable size of 28 to 35 kb (*data not shown*) (**Figure 3C**, **D** and **E**). A fully assembled vector plasmid is then analyzed for sequence and function. The entire engineering process has been streamlined so that new vector genomes can be assembled in about four weeks (*see below*).

**Figure 3:**
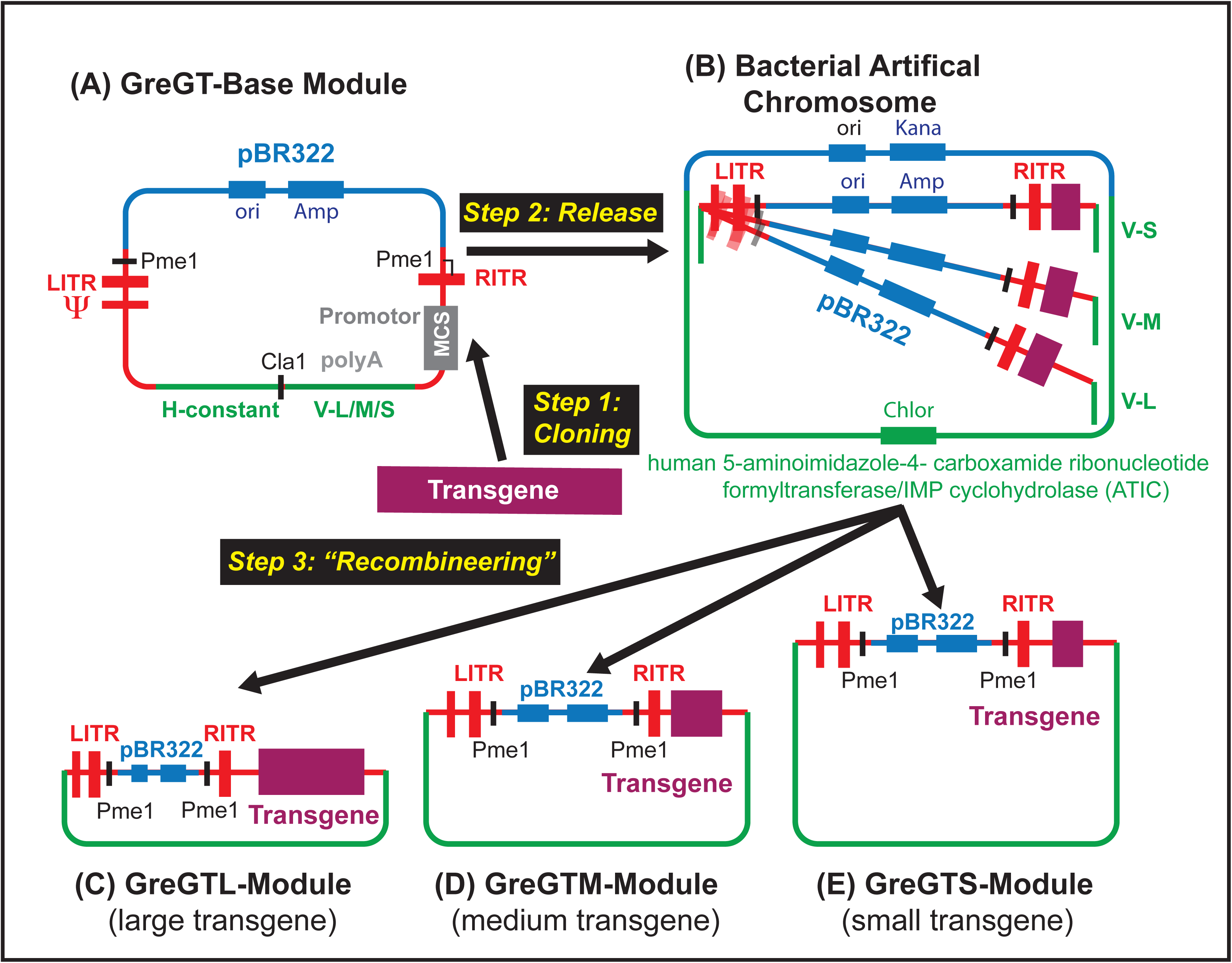
The Design of the Size-Compensating GreGT Vector Genome Modules: A set of three GreGT vector modules (A) assembled in a modified pBR322 plasmid are designed to take up payloads the different sizes. Once a transgene has been integrated, a stuffer sequence of the corresponding size is added by bacterial recombination (B) resulting in an Ad vector genome (C, D, E) of an efficiently packageable size that is released from its cloning vector by a restriction enzyme cut.

#### Packaging Expression Plasmids

For the initial proof-of-principle study, vector modules were packaged in capsids of the human serotype Ad5. Subsequently, we have moved to native or pseudotyped capsids of the rare human Ad6 to minimize an immune interference by pre-existing immune responses. The genome of the Ad6-derived pPaC6 circular expression plasmid was assembled stepwise in a modified pBR322 plasmid. It carries the Ad late genes (L1, L2, L3, L4, L5), as well as the early ones, E2 and E4, and the RITR (**Figure 4**). It was deleted of the LITR, as well as the packaging signal 4′, and the E1 and protein IX genes. A partial deletion of its E3 gene improved its packaging efficacy (*data not shown*). In the pseudotyped pPACs36, the fiber gene was re-engineered as a simian Ad36 and human Ad6 chimera. The simian Ad36 was grafted on the human Ad6 stem. This modification was undertaken to enhance an *intra nasal* uptake of the fdAd vector.

**Figure 4:**
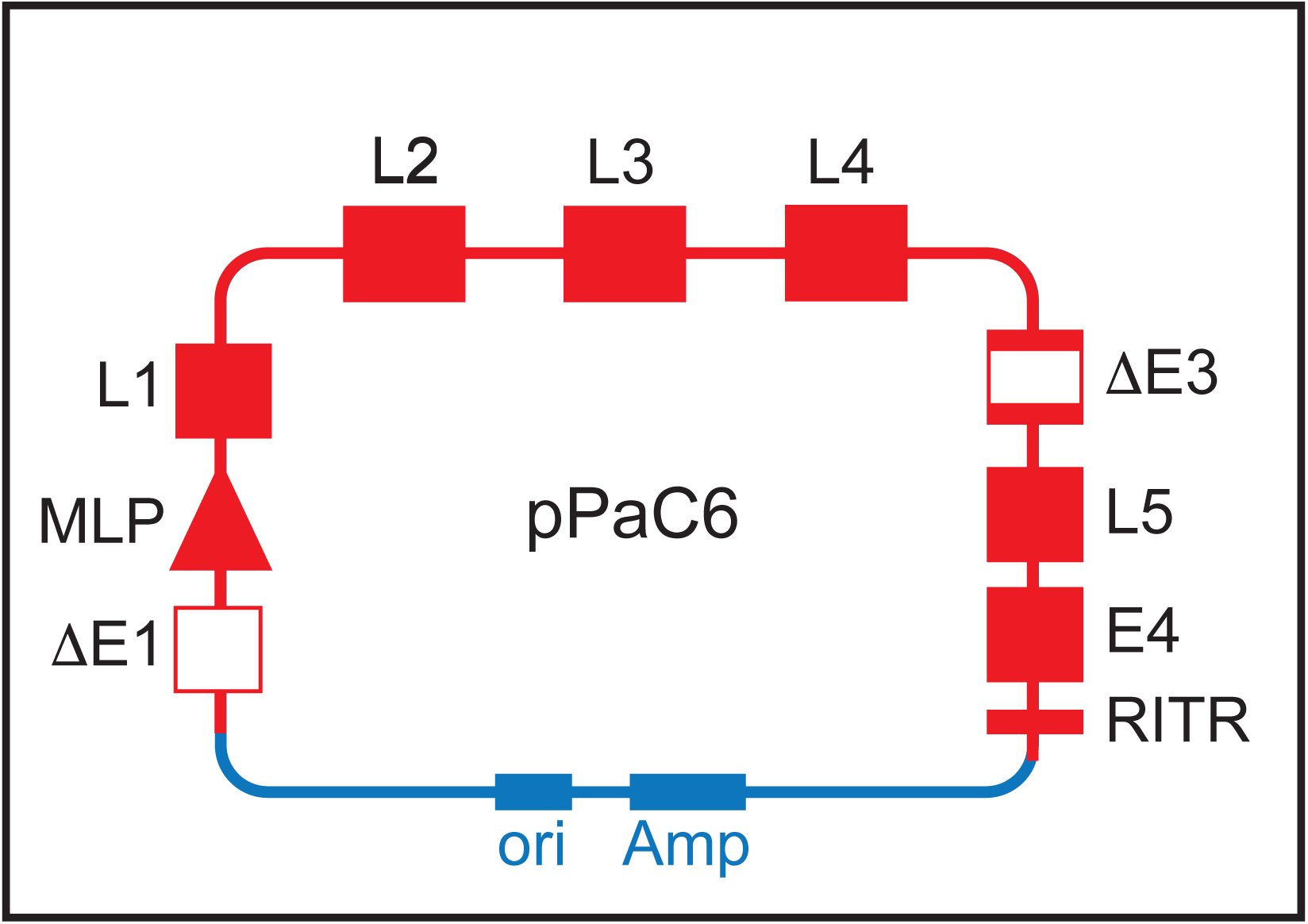
The Construction of a Circular Packaging Expression Plasmid: A model of the circular packaging expression plasmid pPaC6 is presented that is designed to encapsidate fully deleted adenoviral genomes in adenoviral capsids of the human serotype 6.

#### Production Host Cells

Two host cells are commonly used for the production of egAd vectors. First, the HEK293 cell line was developed by a transfection of human embryonic kidney cells with sheared Ad5 DNA (24). The HEK293 cells carry a large Ad genome fragment composed of the Ad E1A and E1B elements, together with the pIX gene (24). Overlaps between the Ad genome in HEK293 cells and minimally modified egAd vectors lead to frequent genetic recombinations resulting in the appearance of RCAs as we and others experienced (25). A new production cell line was established to limit the size of the Ad fragment within the packaging cell. Second, the PER-C6 cells were based on human primary retinal epithelial cells and engineered solely to express Ad E1A and E1B (26). Even with this line of engineered cells, recombinant vector genomes have been reported (27,28). Based on our studies of production rates of of our GreGT vectors in different cell lines, we chose the HEK293 cells as most efficient (*data not shown*). In our system, vector genomes are delivered to the packaging cell as linearized DNA fragments, thus exposing the ITRs to cellular DNases. To protect the integrity of the crucial ITR segments, HEK293 cells were stably transfected with genes for the Ad terminal protein and DNA polymerase. The modified HEK293 cells underwent several rounds of cloning to select the highest producer line (**Figure 5A**). The resulting Q7 cell clone, which delivered an about 100-fold increase in overall production efficiency, was used for all subsequent productions. The engineered Q7 cell retains the gene for the protein IX that stabilizes Ad vectors (29).

**Figure 5:**
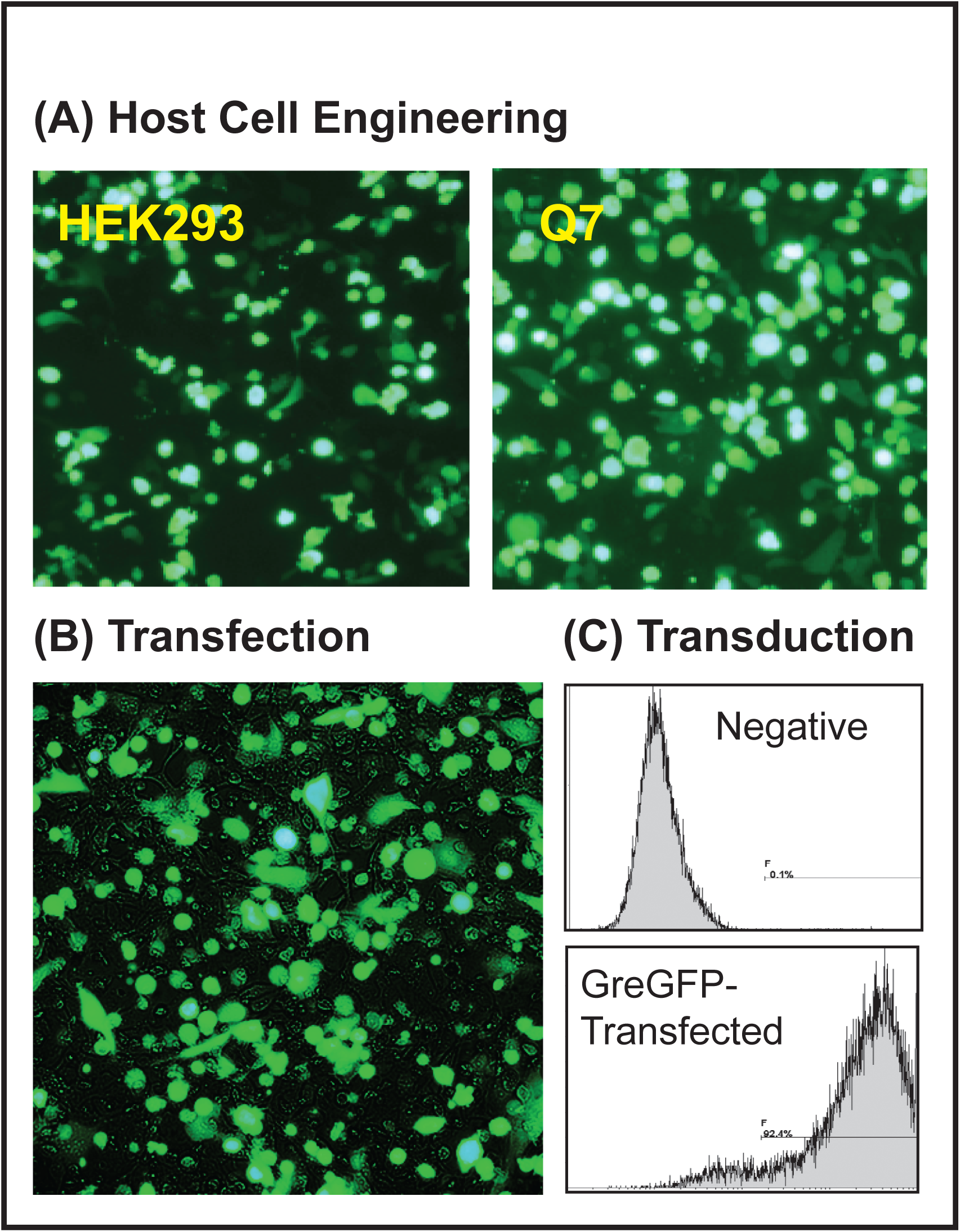
Suveying the Encapsidation Process of a GreGT Vector: (A) A fully deleted adenoviral genome carrying a GFP transgene expression cassette was encapsidated on HEK293 host cells or the variant cell line Q7 that had been stably transfected with the adenoviral terminal protein and DNA polymerase. Samples of vectors produced on either host cell were used to transduce test cells whose GFP expression was examined. (B) Q7 cells co-transfected with a GFP expressing fully deleted adenoviral genome and a circular packaging expression plasmid were examined for GFP expression. (C) The presence of the packaged vector was established by transducing host cells and examining their GFP expression on a FACS.

### Vector Production

#### Vector Encapsidation

Vector production protocols were optimized using the GreGT vector that carried a green fluorescent protein (GFP) gene. In step one, the GreGFP vector genome was released from the cloning plasmid by a simple restriction enzyme cut. The linearized vector carried the packaging signal 4Ψ, the GFP expression cassette, and an appropriate ATIC stuffer fragment, as well as the two ITRs as terminal sequences. It was transfected into Q7 cells together with the circular packaging plasmid, in this case, the Ad5-derived pPAC5. As seen in **Figure 5B**, most Q7 cells lit up green indicating that they had taken up the DNA. After three days of culture, the cells were harvested, the encapsidated vectors were released, purified, and tested for infectivity. As seen in **Figure 5C**, more than 90% of cells expressed GFP when transduced with the GreGFP vector.

After we had been demonstrated that fully deleted vector modules could be efficiently packaged without the use of a helper virus, we investigated whether RCAs could avoided. Aliquots of GreGFP vector preparations were used to infect the non-permissive cell line A549 that lacked Ad genes and therefore could not propagate replication-deficient Ad vectors. These studies indicated that RCAs were not present in preparations of encapsidated GreGFP vectors at levels exceeding 1 in 5 x 10^9^ GE. <u>Vector Purification</u>

Initially, we used a standard ultra-centrifugation protocol to purify encapsidated Ad vectors (30). Although efficient, we believed it impractical for an eventual manufacturing of a clinical product. Therefore, we switched to a two-stage fast protein liquid chromatography system (31) that we adapted to purify our GreGT vectors. The major modification was a switch to a multimodal column for the vector polishing step. When measuring the concentration as active GreGT vectors, the overall recovery through the purification process was in the range of 60 to 80%. We have reached purity levels of vector products that make them suitable to the clinic (*data not shown*). GreGT batches were stored at < - 60°C, at which temperature they remained functionally active for several years (*data not shown*).

### Applications

Once the different components of the GreGT platform had been assembled and optimized, the engineering of a new vector genome should have become straightforward and fast. We asked how long it would take to engineer a new vaccine candidate. The speed of the system was challenged once a new influenza strain had appeared in Asia. In February and March of 2013, three patients were hospitalized in China with a novel lethal avian H7N9 influenza A. On March 31, 2013, the genomic sequences of this influenza virus became publicly accessible. On April 3, an H7N9 hemagglutinin-neuraminidase monocistronic expression cassette was designed and the synthesis of the corresponding DNA fragment was ordered. On April 19, assembly of the vector module was initiated and was completed ten days later by sequence analysis and limited functional testing (**Figure 6**). Subsequent animal trials proved its highly protective function (*data not shown*).

**Figure 6:**
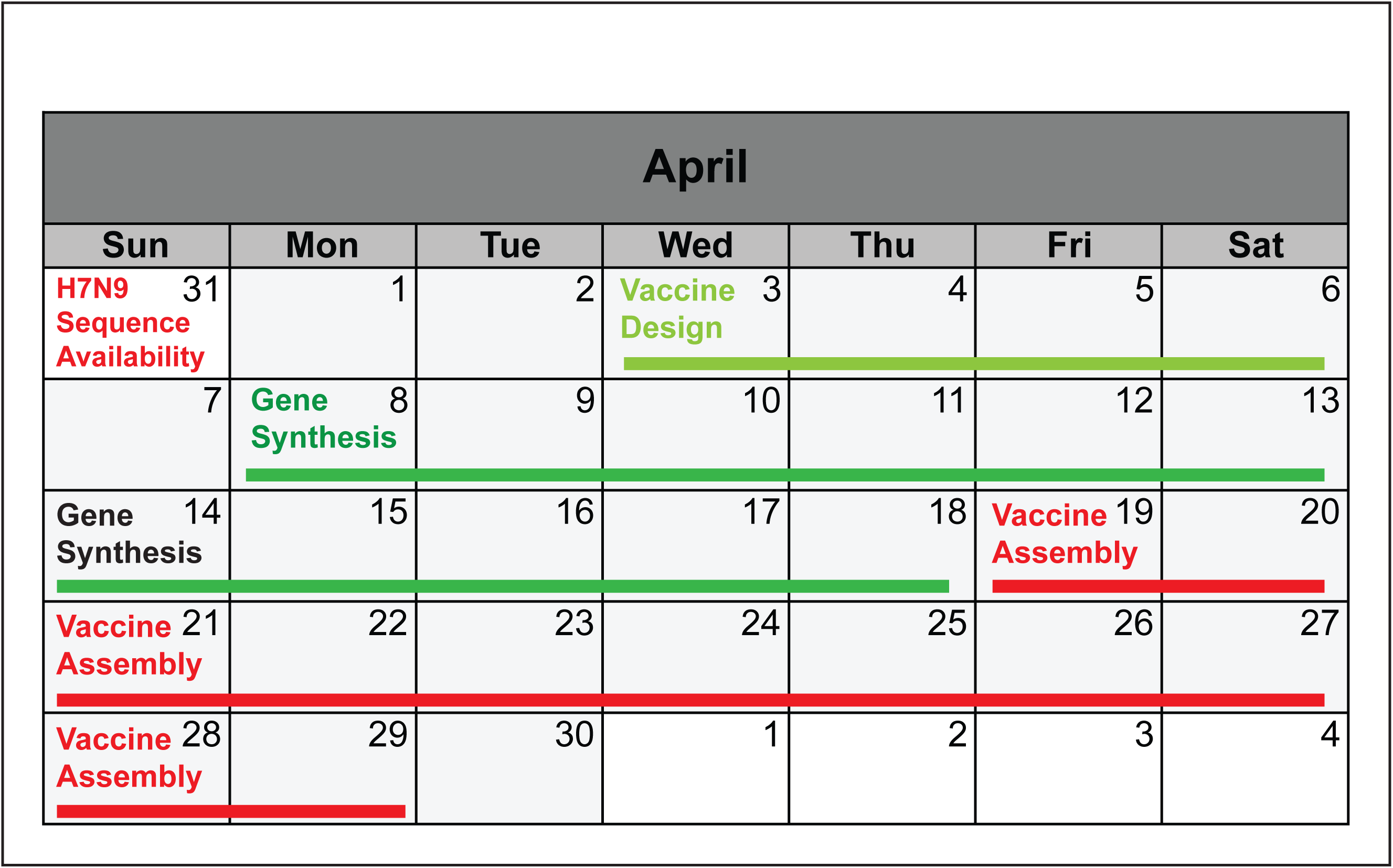
The Timing of the Construction of a Fully Deleted Adenoviral Vector: A calendar excerpt depicts the schedule of the assembly of a GreGT-based H7N9 avian influenza vaccine. The engineering steps are listed as: H7N9 Sequence availability; Vaccine Design; Gene Synthesis; Vaccine Assembly.

#### Flexibility

The production of GreGT vectors is based on two independently modifiable genetic constructs, *i.e.* the vector genome plasmid and the packaging expression plasmid. Therefore, we predicted that exchanging packaging plasmids would lead to the encapsidation of vector genomes into different Ad capsids. A genome, here a vaccine genome targeting infections with the avian influenza virus A/Vietnam/1203/2004 (GreVie), was successfully encapsidated into Ad capsids of the common Ad5 or the rare Ad6 human serotype simply by switching from the pPaC5 to the pPaC6 packaging plasmid (**Figure 7A** and **C**). Others had found that the infection target of an Ad virus or vector depended on the composition of the capsid fiber (32). We therefore investigated whether the pPaC6 packaging plasmid could be pseudotyped with a chimeric fiber gene. The knob of the simian Ad36 was grafted on the stem of the human Ad6. As seen in **Figure 7**, the pseudotyped packaging plasmid was still able to infect the test cells. These experiments demonstrated the inherent flexibility of the GreGT technology. They showed that an Ad capsid could be replaced by simply swapping the packaging plasmid during vector production.

**Figure 7:**
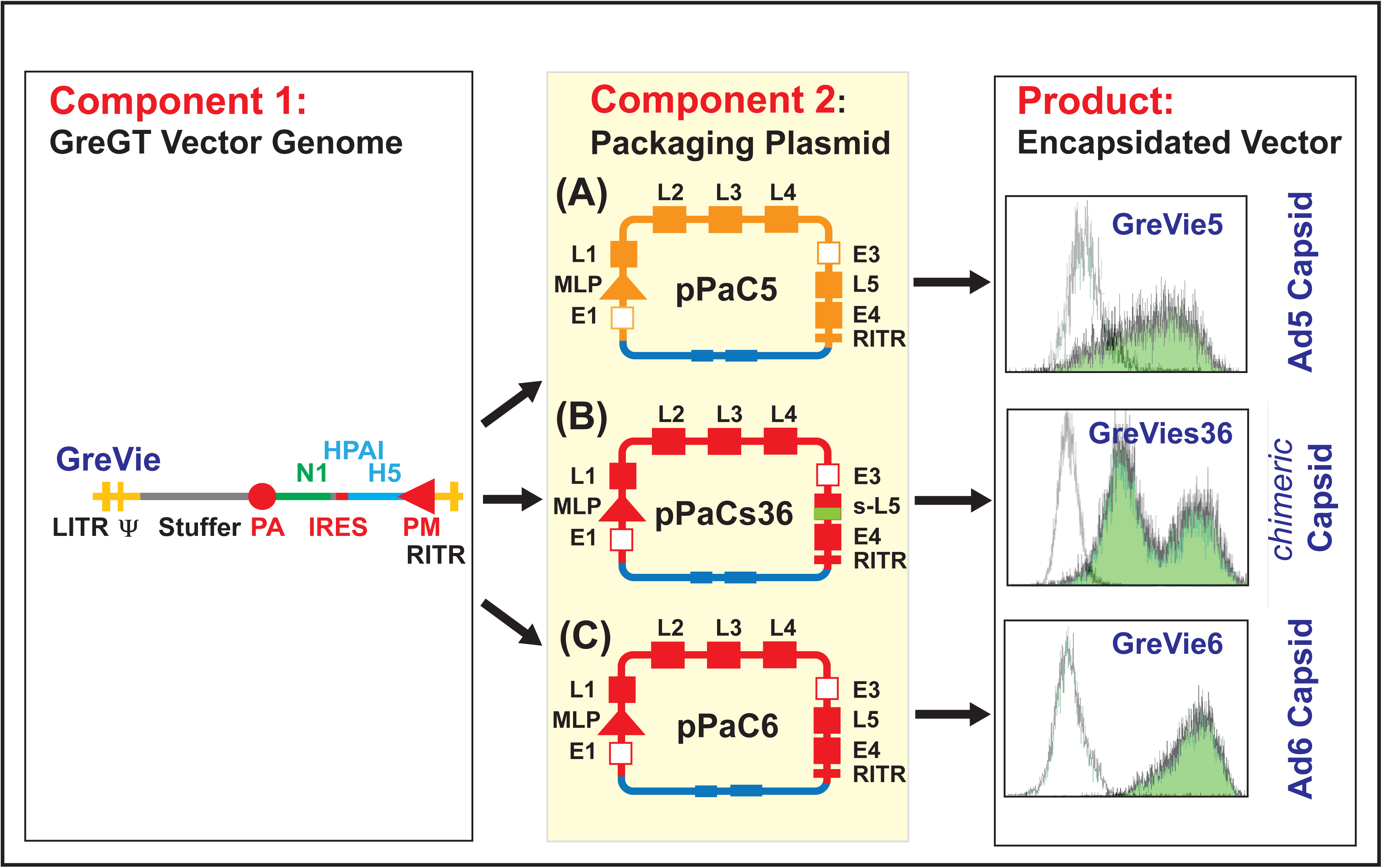
The Flexibility of the GreGT Vector Platform: The flexibility of the two-component GreGT *plug-and-play* technology is demonstrated by the packaging a fully deleted adenoviral genome (GreVie) of an avian influenza vaccine (Component 1) with the help of circular packaging expression plasmids (Component 2) of the of human adenoviral serotypes 6 (pPaC6) and 5 (pPaC5) and of a fiber pseudotyped adenoviral capsid (pPaC6s36). The different encapsidated vectors (Product), *i.e.* GreVie5, GreVies36 and GreVie6, were used to transduce test cells, on which the surface expression of the hemagglutinin transgene was determined by immunofluorescence.

#### Versatility

The GreGT platform was designed to address a broad spectrum of gene transfer-based human therapies. Exchanging the vector genome construct during vector production should be able to be used to address new applications. **Figure 8A** presents three different examples. As a vaccine model, the GreVie vector genome was used (*see above*). Its packaging into an Ad6 capsid was compared to that of the tissue engineering vector genome GreCD8 that carries the mouse CD8 α-chain gene (**Figure 8B**). This example was chosen, as we had previously determined in animal studies that pancreatic islets transduced to express the CD8 α-chain were protected from rejection when grafted into fully allogeneic recipients (17). As a third example of the breadth of the GreGT platform, a gene therapy vector was assembled. Here, the large capacity (up to 33 kb) of the fully deleted vector was utilized when a transgene expression cassette carrying the complete human hemagglutination factor VIII cDNA (7 kb in length) was loaded (**Figure 8C**). Even larger genes, such as the the USH2A gene of about 15 kb in length could easily be accommodated in a GreGT vector (*data not shown*). In addition to these examples, we have examined whether the available space would allow the transport of arrays of genes in a single vector, such as several vaccine antigen genes targeting the same or different infections or entire enzyme cascades for gene therapy.

**Figure 8:**
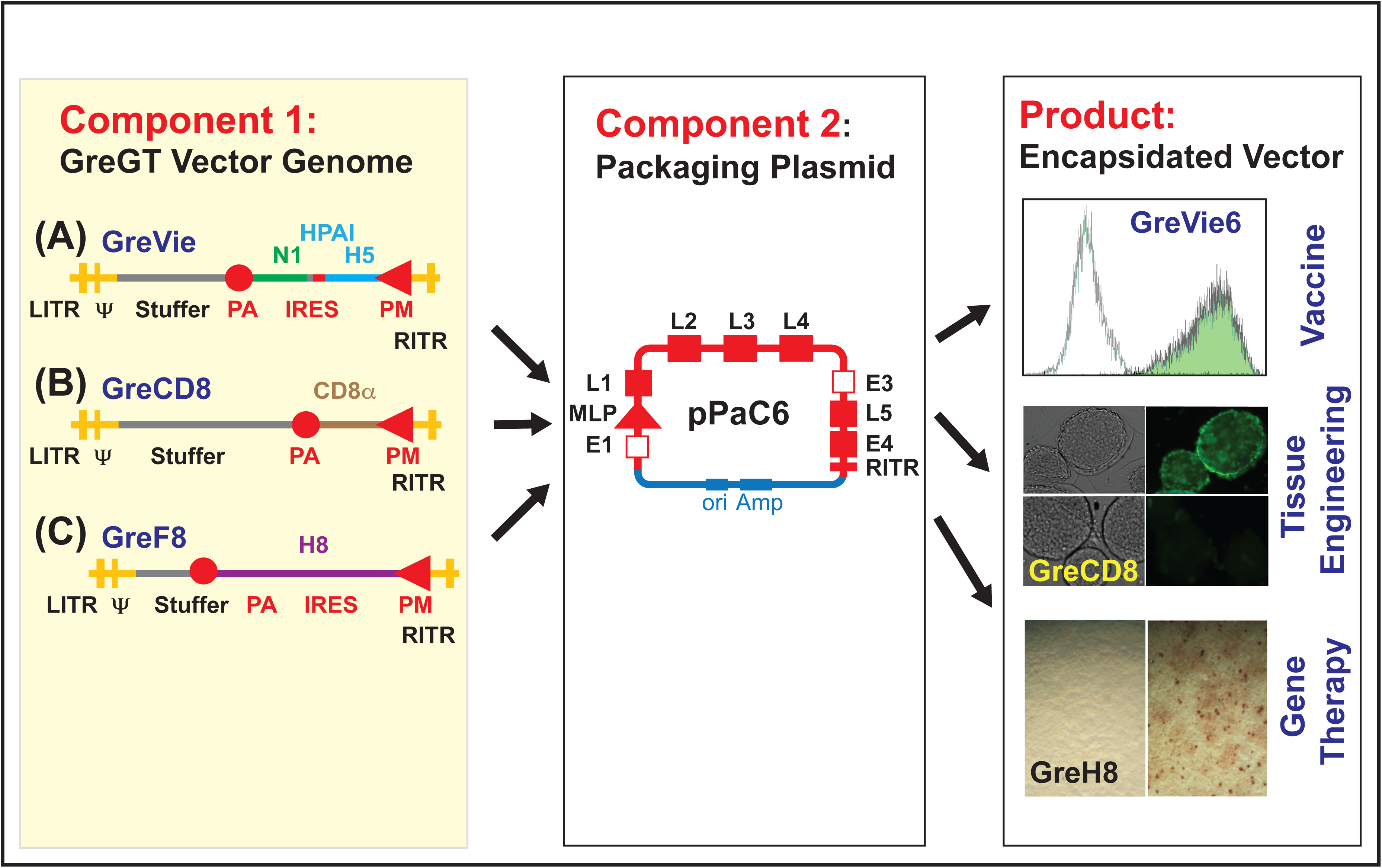
The Versatility of the GreGT Vector Platform: The versatility of the two-component GreGT *plug-and-play* technology is demonstrated by the packaging of fully deleted adenoviral genomes (Component 1) of an avian influenza vaccine (GreVie), an immune inhibition vector (GreCD8) or a coagulation factor VIII genetransfer vector (GreF8) with the help of the circular packaging expression plasmi (Component 2) of the of human adenoviral serotypes 6 (pPaC6) and 5 (pPaC5). The different encapsidated vectors (Product) were used to transduce test cells, on which the expressions of the hemagglutinin transgene (GreVie6) and the CD8 (GreCD8) were determined by immunofluorescence and the one of the coagulation factor VIII (GreF8) by an ELISpot assay.

#### Immunogenicity

Immune responses to the endogenous Ad gene components encoded in the genome of egAd or Ad helper viruses may interference with the gene transfer success. An important goal of the creation of the GreGT platform was to minimize such immune responses. Mice immunized *i.m.* with a single dose of an egAd raised strong immune responses as depicted in **Figure 1**. Whe we examined the immunogenicity of the GreGT-based avian influenza vaccine GreVie vaccine, we saw a different result. Mice that had received a single *i.m.* injection did not mount significant anti-Ad humoral immune responses (**Figure 9**). Yet, strong serum antibody levels were raised against the hemagglutinin antigen encoded within the GreVie transgene expression cassette. These data demonstrated that our design goal of reduced vector immunogenicity was indeed met.

**Figure 9:**
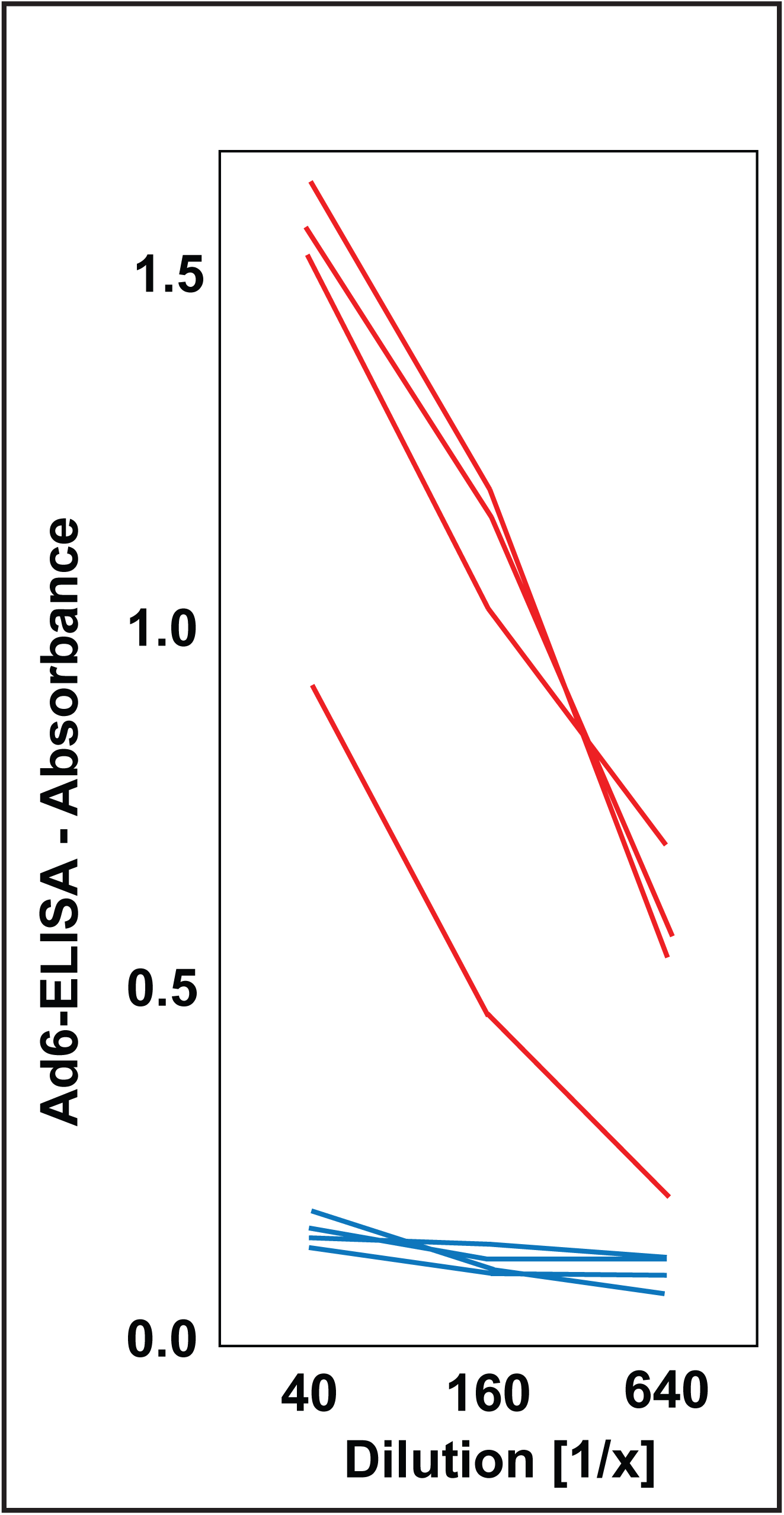
Minimizing the Immunogenicity of the Vector Carrier: Sera of mice that had be immunized by an *intra muscular* injection with the GreGT-based avian influenza vaccine (GreVie6) were tested for humoral immune responses against the adenoviral capsid of the human serotype 6 (blue lines) or the hemagglutinin transgene (red lines).

### Summary

We have developed a new gene transfer platform that allowed the production of fully deleted Ad vectors without the participation of a helper virus constructs. It prevents the presence of helper viruses or RCAs in the therapeutic product. It provides great flexibility and versatility, as it allows easy exchanges of the Ad capsids and the packaged genomes during vector production. The reduced immunogenicity to its vector carrier should minimize the interference by both pre-existing and induced immune responses. Repeated uses of the same vector should be facilitated by limiting the deleterious effects of vector-directed immune responses. The large payload permits the delivery of large genes, as well as arrays of gene that cannot be delivered by other existing vector systems. The vector construction process has been streamlined. The speed of the present vector platform in vector design and standardized production may open the door to a new paradigm of vaccination. Rather than preemptively storing vaccines aimed at the most probable outbreaks, it may be possible to specifically and swiftly design vaccines targeting newly emerging infections with the help of our optimized plug-and-play GreGT platform.

## MATERIAL AND METHODS

The **fdhiAd base modules** (GreS, GreM and GreL) were constructed in the pBR322 plasmid (ATCC) with fragments derived from a human Ad of the serotype 5 (GenBank Accession Number AC00008) (**Figure 3**). After a multiple cloning site (MCS) had been ligated into the pBR322 Sal1/EcoR1 site destroying the tetracycline resistance gene, the following components were stepwise inserted: the Ad 5’ left inverted terminal repeat (LITR) - 4′- segment with an attached Pme1 site; a region (H-constant) homologous to exon 3 of the human 5-aminoimidazole-4-carboxamide ribonucleotide formyltransferase (ATIC; GenBank Accession Number NG013002) gene (for GreS, GreM, GreL); a Cla1 site; one of three regions (V-S, V-M, and V-L) homologous to different segments of the human ATIC gene (nucleotides 73,026 through 73,385, nucleotides 78,516 through 78,875, nucleotides 83,616 through 84,095) (for GreS, GreM, GreL); the human growth hormone polyadenylation signal sequence (HGH); the cytomegalovirus (CMV) immediate/early promoter; and the Ad 3’ right inverted terminal repeat (RITR) segment with a Pme1 site. Upon integration of a transgene*, stuffer* DNA was added from a bacterial artificial chromosome (BAC) carrying exons 3 through 16 of the human ATIC, into which a chloramphenicol resistance marker had been moved (BAC-ATIC, D. Paterson, University of Denver, Denver, CO). For this purpose, a transgene-carrying base vector construct was transformed into bacteria (D10β) carrying the BAC-ATIC as well as the λ phage recombination protein expressing pRED/ET plasmid (GeneBridges). With homologies of the H-constant region of the fdhiAd vector constructs to ATIC exon 3 and those of the V-S, V-M, or V-L regions, stuffers of about 15 kb, 10 kb, and 5 kb were added. The chloramphenicol marker was excised. The *Aequorea victoria* green fluorescent protein (Life Technologies) or the influenza antigen monocistronic construct of codon optimized hemagglutinin and neuraminidase (A/Vietnam/1203/2004), separated by an encephalomyelitis virus internal ribosome entry site (IRES) (Clontech), were cloned into the MCS of an GreS base vector. Transgene cassettes for the coagulation factor VIII and the USH2A genes were moved into the GreM base vector modules.

The **engineering of the packaging construct pPAC6** (**Figure 4**) was assembled as follows. A DNA fragment composed of a poly-linker (EcoRI - AflII - AvrII) followed by the Ad sequence bp 35,289 to 35,759 and a BamHI site were directly cloned into the EcoRI - BamHI site of the pBR322 plasmid. An AflII - AvrII fragment of the wild-type Ad6 genome (bp 3,549 to 13,068) was directly cloned into the respective sites of the modified pBR322 plasmid. This plasmid was linearized using AvrII. It was *recombineered* with a ClaI fragment bp (917 to 35,769) of the wild-type Ad6 genome with the help of a bacterial recombinase expressed by the pRed/ET vector (*GeneBridges*). This added the Ad6 sequences bp 13,068 to 35,289 to the plasmid. A kanamycin-resistance sequence (*Sigma-Aldrich*) was centered between a *left* Ad6 homology arm (bp 29,290 to 29,340) and a *right* Ad6 homology arm (bp 30,146 to 30,195). This DNA fragment was also moved into the packaging construct by *recombineering*. This replaced Ad6 sequences bp 29,341 to 30,145 with a kanamycin resistance gene deleting the activities of Ad6 E3, 13.4k, E3 gp 19k, E3 CR1 beta, E3 RID alpha, E3 RID beta and E3 14.7k .

The cloned **packaging cell line Q7** was derived from a human embryonic kidney cell line (HEK293, ATCC). The Q7 cells were stably transfected with a TP/Pol expression vector that, based on pUC57 (GeneScript), carried a synthesized DNA cassette containing an *artificial* polyadenylation sites, the human Ad5 terminal protein, the human surfeit bidirectional promoter (GenBank Accession Number AC002107), the human Ad5 DNA polymerase, and the human herpes simplex virus 5 puromycin resistance marker (Clontech).

To **encapsidate** GreGT vector genomes, adherent Q7 cells were expanded in tissue culture flasks (Greiner) to about 90% confluency in Iscove’s modified Eagle’s medium (IMDM, Gibco) supplemented with 10% fetal bovine serum (Life Technologies) (33).

GreVac vector module and packaging construct DNAs mixed with polyethyleneimine (PolyPlus) were added to the Q7 cells. After 72 hours the transfected Q7 cells were harvested with Trypsin (Sigma) in PBS, spun down and resuspended in virus release storage medium.^i^ They underwent three rounds of freezing and thawing to release the encapsidated vectors. After removal of cellular debris by centrifugation, the vector particles were purified by two-step column chromatography. The clarified vector suspension is loaded onto an anion exchange column, Capto Q (*Cytiva*), equilibrated to 150 mM NaCl, and eluted by a NaCl gradient to 600mM. The same buffer is used with modified NaCl concentrations: Tris-HCl 20 mM pH 7.8, MgCl_2_ 1 mM, EDTA 0.1 mM, Sucrose 2% (W/V) and polyethylene glycol (PEG) 4000 0.002mM. The active fractions are pooled and *polished* on a multi-model column, Capto Core 700 (*Cytiva*), with a buffer consisting of Tris-HCl 20 mM pH 7.8, NaCl 450 mM, MgCl_2_ 1 mM, EDTA 0.1 mM, Sucrose 2% (W/V) and polyethylene glycol (PEG) 4000 0.002mM. The vector suspension is concentrated on a small Capto Q (*Cytiva*) column and then adjusted to the final concentration of the vaccine product of 1 x 10^9^ GE/ml in the Vector Formulation Buffer (VFB) consisting of Tris-HCl 20 mM pH 7.4, NaCl 150 mM, MgCl2 1 mM, EDTA 0.1 mM, Sucrose 2% (W/V) and polyethylene glycol (PEG) 4000 0.002mM. The final vaccine products are filter-sterilized prior to vialing. Their concentrations are measured as infectious particles by expression of the transgene or as genome equivalents by quantitative PCR using polynucleotide pairs targeting the transgene expression cassette.

To detect **RCA**s, samples of purified vectors were added to cultured A549 cells (ATCC) at increasing numbers (between 1 x 10^6^ and 5 x 10^10^ GE). Samples (between 2 and 1 x 10^3^ plaque forming units) of defined wild-type human Ad serotype 5 or 6 preparations (ATCC) were used to standardize the assay. After 48 hours of culture, the potential viruses were harvested and added to secondary cultures of A549 cells. After another 48 hours the cells were fixed in cold methanol (Sigma) and stained with an anti-Ad hexon antibody (Sigma), whose binding was revealed with a horseradish peroxidase labeled rabbit anti-mouse antibody (Southern Biotechnology). The plates were scored by conventional microscopy. The sensitivity of the assay was determined as 5 PFU of the wild-type Ad virus in a test preparation.

The **functional activity** of the different vector constructs was measured by adding graded amounts of a purified vector to Q7 cells. GFP expression was directly measured on a Cytomics FC500 (Beckman-Coulter) or a fluorescence microscope (Eclipse 80i, Nikon). Surface expression of influenza hemagglutinin was detected with an inhouse-produced polyclonal anti-hemagglutinin antibody preparation, for which goats had been hyper- immunized with the purified H5 influenza protein (Protein Sciences). The extent of anti-hemagglutinin antibody binding was revealed with a fluoresceinated rabbit anti-goat antibody (Southern Biotechnology). Expression of the mouse CD8 α-chain was determined with fluoresceinated antibodies (Thermofisher). Secretion of the human coagulation factor VIII was determined by EliSpot assays, in which the factor was captured and developed by specific antibodies linked to horseradish peroxidase (Invitrogen).

Balb/c were **immunized** *i.m.* with 1 x 10^8^ genome equivalents (GE) of the egAd(empty) or the GreVie vaccine candidate. Twenty-eight (28) days thereafter, the presence of anti-Ad and anti-hemagglutinin antibodies was measured by ELISA against a purified Ad6 virus preparation (Greffex, Inc.) and a recombinant hemagglutinin protein (Creative Biomart). In the first case, U-bottomed 96-well plates (Corning) were coated with an anti-Ad antibody (Creative Biolabs). After Ad6 viruses had been added and the plates had been blocked with bovine serum albumin (Sigma), dilutions of the mouse sera were added. In the second case, the plates were coated with the hemagglutinin protein, then blocked. Then, dilutions of the mouse sera were added. Binding of the mouse antibodies were detected by a rabbit anti-mouse antibody labeled with horseradish peroxidase (Southern Biotechnology). An in-house produced goat anti-hemagglutinin monoclonal antibody was used as positive control standard.

## ACKNOWLEDGEMENTS

This research was supported in part by cooperative agreements and grants from the National Institute of Standards and Technology and the National Institutes of Health.

